# In silico identification and validation of inhibitors of the interaction between neuropilin receptor 1 and SARS-CoV-2 Spike protein

**DOI:** 10.1101/2020.09.22.308783

**Authors:** Samantha Perez-Miller, Marcel Patek, Aubin Moutal, Carly R. Cabel, Curtis A. Thorne, Samuel K. Campos, Rajesh Khanna

## Abstract

Neuropilin-1 (NRP-1) is a multifunctional transmembrane receptor for ligands that affect developmental axonal growth and angiogenesis. In addition to a role in cancer, NRP-1 is a reported entry point for several viruses, including severe acute respiratory syndrome coronavirus 2 (SARS-CoV-2), the causal agent of coronavirus disease 2019 (COVID-19). The furin cleavage product of SARS-CoV-2 Spike protein takes advantage of the vascular endothelial growth factor A (VEGF-A) binding site on NRP-1 which accommodates a polybasic stretch ending in a C-terminal arginine. This site has long been a focus of drug discovery efforts for cancer therapeutics. We recently showed that interruption of the VEGF-A/NRP-1 signaling pathway ameliorates neuropathic pain and hypothesize that interference of this pathway by SARS-CoV-2 spike protein interferes with pain signaling. Here, we report hits from a small molecule and natural product screen of nearly 0.5 million compounds targeting the VEGF-A binding site on NRP-1. We identified nine chemical series with lead- or drug-like physico-chemical properties. Using an ELISA, we demonstrate that six compounds disrupt VEGF-A-NRP-1 binding more effectively than EG00229, a known NRP-1 inhibitor. Secondary validation in cells revealed that almost all tested compounds inhibited VEGF-A triggered VEGFR2 phosphorylation. Two compounds displayed robust inhibition of a recombinant vesicular stomatitis virus protein that utilizes the SARS-CoV-2 Spike for entry and fusion. These compounds represent a first step in a renewed effort to develop small molecule inhibitors of the VEGF-A/NRP-1 signaling for the treatment of neuropathic pain and cancer with the added potential of inhibiting SARS-CoV-2 virus entry.

## Introduction

As of September 21, 2020 COVID-19 has infected more than 31 million people and caused nearly 1 million deaths worldwide^1^. This disease is caused by severe acute respiratory syndrome coronavirus 2 (SARS-CoV-2) which primarily gains entry to cells via binding of SARS-CoV-2 Spike glycoprotein to angiotensin converting enzyme 2 (ACE-2) and subsequent endocytosis^2-4^. Recent reports have identified additional entry points, including neuropilin 1 (NRP-1) ^5, 6^.

Neuropilins are cell surface receptors for secreted glycoproteins with roles in neural outgrowth, cardiovascular development, immune response, as well as tumor growth and vascularization^7, 8^. Two neuropilin isoforms, NRP-1 and NRP-2, share ∼44% sequence identity in humans and function in different pathways^7, 8^. Both share a modular architecture with three extracellular domains, a single transmembrane helix, and a short cytoplasmic tail (**Figure 1A**)^9^. The a1a2 modules, homologous to CUB (for complement C1r/C1s, Uegf, Bmp1) domains, interact with semaphorin 3A (SEMA3A) to mediate stimulation of growth cone collapse in developing neurons^7^. The b1b2 modules are homologous to the C-terminal domains of blood coagulation Factors V and VIII^7^. The c domain, homologous to meprin, A5, and mu-phosphatase (MAM), was initially thought to be involved in dimerization, but is more likely to contribute to complex assembly by positioning the preceding domains away from the membrane^10^. The single transmembrane helix contributes to homo and heterodimerization^11^ and the C-terminal cytoplasmic tail is thought to contribute to signaling through synectin, stimulating receptor-mediated endocytosis^8^.

**Figure 1.**
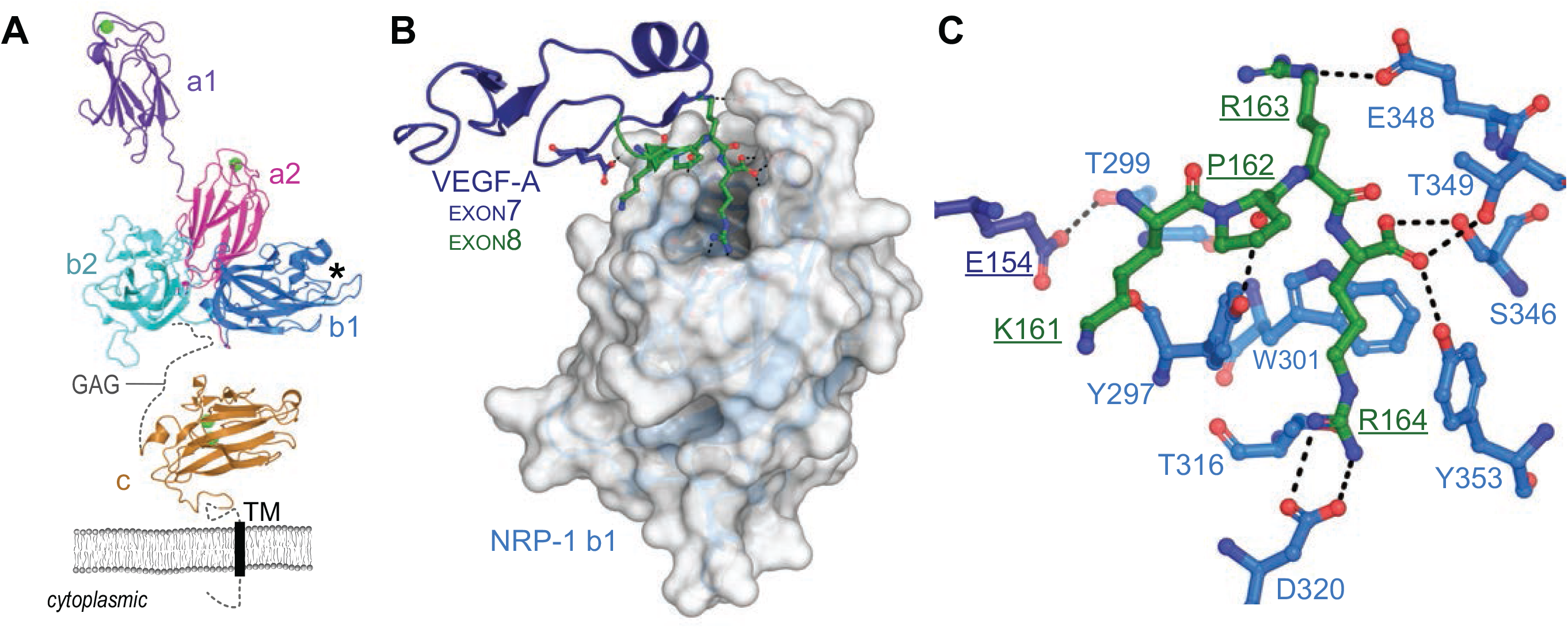
Schematics of NRP-1 domains, VEGF-A_165_ binding, and CendR interaction network. **A**. Domain architecture of NRP-1 with domain a1 from mouse (PDB ID 4gz9)^78^, domains a2, b1, b2 (PDB ID 2qqm)^9^ and MAM (PDB ID 5L73)^10^ from human. Bound Ca^2+^ shown as green spheres, missing loops as dashes, transmembrane domain as a rectangle. GAG indicates region of glycosaminoglycan modification. *Indicates VEGF-A interaction pocket. **B**. Structure of NRP-1 b1 domain, shown as white surface, in complex with the heparin binding domain (exon 7/8) of VEGF-A_164_, shown as cartoon with exon 7 in dark blue, exon 8 in green (PDB ID 4deq)^14^. **C**. Details of VEGF-A Glu154 and KPRR_164_ interactions with NRP-1 (close up of view B). Dashes indicate polar or salt-bridge contacts within 3.0 Å.

It is through the b1 module that SARS-CoV-2 may gain entry, by taking advantage of the interaction site for vascular endothelial growth factor A (VEGF-A). VEGF-A isoforms and other growth factors that terminate in a polybasic stretch ending with an obligatory arginine residue, termed the C-terminal end arginine (CendR) rule^12^, interact with an acidic pocket formed by loops extending from the beta-barrel of b1 (**Figure 1B**)^13, 14^. The structure of the heparin binding domain of the 164 residue isoform of VEGF-A (VEGF-A_164_) confirmed that the C-terminal Arg (Arg164) engages the NRP-1 b1 domain pocket with the guanidine forming a bidentate salt-bridge with conserved Asp320 and the carboxylate forming hydrogen bonds to conserved Ser346, Thr349, and Tyr353 (**Figure 1C**)^14^. It is notable that VEGF-A binds to both NRP-1 and NRP-2, but has higher affinity for NRP-1 due to amino acid substitutions within the first loop region of NRP-1 that provide additional contacts between NRP-1 Thr299 and VEGF-A Glu154 (**Figure 1B, C**)^14^. Furthermore, although b1 and b2 are homologous domains, critical residues within the loops of the binding domain are not conserved with the result that the CendR interaction site is not present on b2.

VEGF-A has long been known for its essential role in blood vessel growth and function, but has more recently been shown to be pro-nociceptive^15^. VEGF-A is a selective endothelial cell mitogen that promotes angiogenesis, primarily via interaction with the VEGF receptor VEGFR2, also known as kinase insert domain-containing receptor (KDR)^16^. Alternative splicing of the VEGF-A gene produces several isoforms of the mature protein containing between 121 and 206 amino acid residues, with VEGF-A_165_ being pro-nociceptive^17^ via sensitization of transient receptor potential (TRP) channels^18^ and ATP-gated purinergic P2X_2/3_ receptors^19^ on dorsal root ganglion (DRG) neurons. This alternative splicing is dependent on serine-arginine rich protein kinase 1 (SRPK1) which mediates the phosphorylation of serine-arginine rich splice factor (SRSF1)^17, 20, 21^. VEGF binding to VEGFR2, a co-receptor for NRP-1, is associated with receptor dimerization and activation that triggers downstream signaling pathways including phosphatidylinositol 3-kinase (PI3-K)/Akt and phospholipase C gamma/extracellular signal-regulated kinase (PLCg/ERK)^16^. Clinical findings that VEGF-A contributes to pain are supported by observations that in osteoarthritis increased VEGF expression in synovial fluids has been associated with higher pain scores^22^. VEGF-A has been reported to enhance pain behaviors in normal, nerve-injured and diabetic animals^17, 23^.

It is known that neuropilins are entry points for several viruses, including human T-lymphotropic virus-1 (HTLV-1)^24^ and Epstein-Barr virus (EBV)^25^. In both cases, furin processing of viral glycoproteins results in polybasic CendR motifs that directly interact with the VEGF-A site on the NRP-1 b1 domain^26^. Compared to SARS-CoV-1, the causative agent of SARS, mutation in the furin cleavage site of SARS-CoV-2 results in production of a CendR motif (^682^RRAR^685^) which was shown to bind to the neuropilin b1 VEGF-A site, suggesting NRP-1 as a possible route of viral entry^5,6^. The importance of NRP-1 is supported by recent evidence of upregulated NRP-1 in lung samples from COVID-19 patients^5^.

These connections raise an interesting question: does interference with the VEGF-A/NRP-1 signaling pathway by SARS-CoV-2 result in dampened pain? This question has been examined by our laboratory; we recently showed that SARS-CoV-2 Spike protein binding to NRP-1 prevents VEGF-A signaling and reduces neuropathic pain in an animal model^27^. Thus, NRP-1 represents a novel target for treating neuropathic pain^27^. Furthermore, targeting NRP-1 also presents a unique approach to inhibiting viral entry and/or re-entry into cells to reduce viral load. Due to its role in cancer, NRP-1 has been a target for drug design for over 20 years. During this time, discovery efforts have focused on development of NRP-1 antibody therapies^9, 28-33^, including a recent dual-specificity antibody to VEGFA and NRP-1^32, 33^, peptides that target transmembrane domain interactions^11, 34-37^ or the CendR interaction site^12, 31, 38-53^, as well as small-molecules that target the CendR site ^54-61^. With one exception^43^, the CendR site targeting peptides contain a CendR motif, even those which are cyclical^39, 45, 46, 49^ or branched^51, 53^. Two of the most well-known small molecule NRP-1 inhibitors, EG00229 and EG01377, contain a terminal Arg-like moiety and carboxyl group known to be key to interaction with NRP-1^54, 61^ as does the newly developed fluorescent compound based on EG01377^62^. The remaining known small molecule compounds consist of an arginine-derivative series^60^ and several diverse chemotypes such as acylthioureas^56^, benzamidosulfonamides^59^, bis-guanidines^55^, or aryl benzylethers^58^. Many of these small molecule compounds contain functional groups of suboptimal physico-chemical and ADME properties.

In order to identify unique compounds that could be used to interrogate the role of these signaling complexes in pain, we conducted a virtual screen of nearly 0.5 million compounds against the NRP-1 CendR site, resulting in nearly 1,000 hits. Here, we present 9 chemical series of synthetic and natural compounds with lead- or drug-like physico-chemical properties and identify a pharmacophore model which will guide future design of compounds. We also show that six of the compounds interfere with VEGF-A binding more effectively than EG00229, a known NRP-1 inhibitor and three interfere with VEGF-A binding as well as the furin-cleaved Spike S1 domain. Furthermore, our compounds show inhibition of VEGF-A triggered VEGFR2 phosphorylation in a cell-based assay. Finally, two of the compounds inhibited antiviral activity.

## Results and Discussion

We conducted virtual screens against the VEGF-A binding site on the NRP-1 b1 domain using three libraries: a ∼211 K synthetic compound library (DIV) from ChemBridge; a ∼257 K natural compound library (NC1) obtained from the COlleCtion of Open NatUral producTs (COCONUT) resource^63^; and a ∼20 K (NC2) natural compound library from the ZINC15 database^64^. The screens were run once without ligand-receptor interaction constraints and repeated with the constraint that compounds form a hydrogen bond to Asp 320, a key residue for coordinating the terminal arginine in the CendR motif^14^. This constraint was used in attempt to select for compounds that interact in a similar way as observed for VEGF-A^14^, known inhibitors^42, 54, 60, 61^ and modeled SARS-CoV-219 CendR terminal arginine^6^. However, application of this constraint led to reduced overall scores and strained conformations for most compounds in the DIV and NC1 libraries. Therefore, we report only the unconstrained screen results for these libraries and both constrained and unconstrained results for the NC2 screen.

### Selection of top compounds

The combined output of the four screens produced a total of 1,147 hits. Compounds from each screen were sorted by Glide XP GScore (kcal/mol) and visually inspected for substructure match of core scaffolds and patterns of chemically reactive moieties while considering the diversity of chemotypes. Compound representatives with scaffold decoration that suggested initial structure activity relationships (SAR) were extracted and grouped into series resulting in a set of nine diverse chemotypes (series). These series include both small synthetic molecules and natural products (**Table 1**). While calculated property ranges vary for the chemotype and drug indication, we intend to keep compounds in our series such that molecular weight (Mw) < 400 Da, calculated octanol-water partition coefficient (clogP) < 3.5, and calculated solubility (clogS) at pH 7.4 > –5 M. Consequently, we expect the initial absorption, distribution, metabolism, and excretion (ADME) profile of the series to be acceptable (e.g., good hepatocyte clearance and bioavailability). To assess the probability that hit series compounds are orally available, we determined components and overall compliance with Lipinski rule of 5 (Ro5)^65, 66^. Since the Ro5 guidelines were derived for orally available small molecules, we did not use this binary parameter for the classification of natural products. Also, since solubility and CNS penetration rules originated from small molecule sets, these calculations are not applicable to natural products and were also omitted. Furthermore, detailed inspection of data of **Table 1** revealed that natural product compounds **11, 12**, and **15** are in fact small molecule chemotypes. Correspondingly, all molecular descriptors were calculated for these compounds and they were considered part of the small molecule set.

**Table 1.**
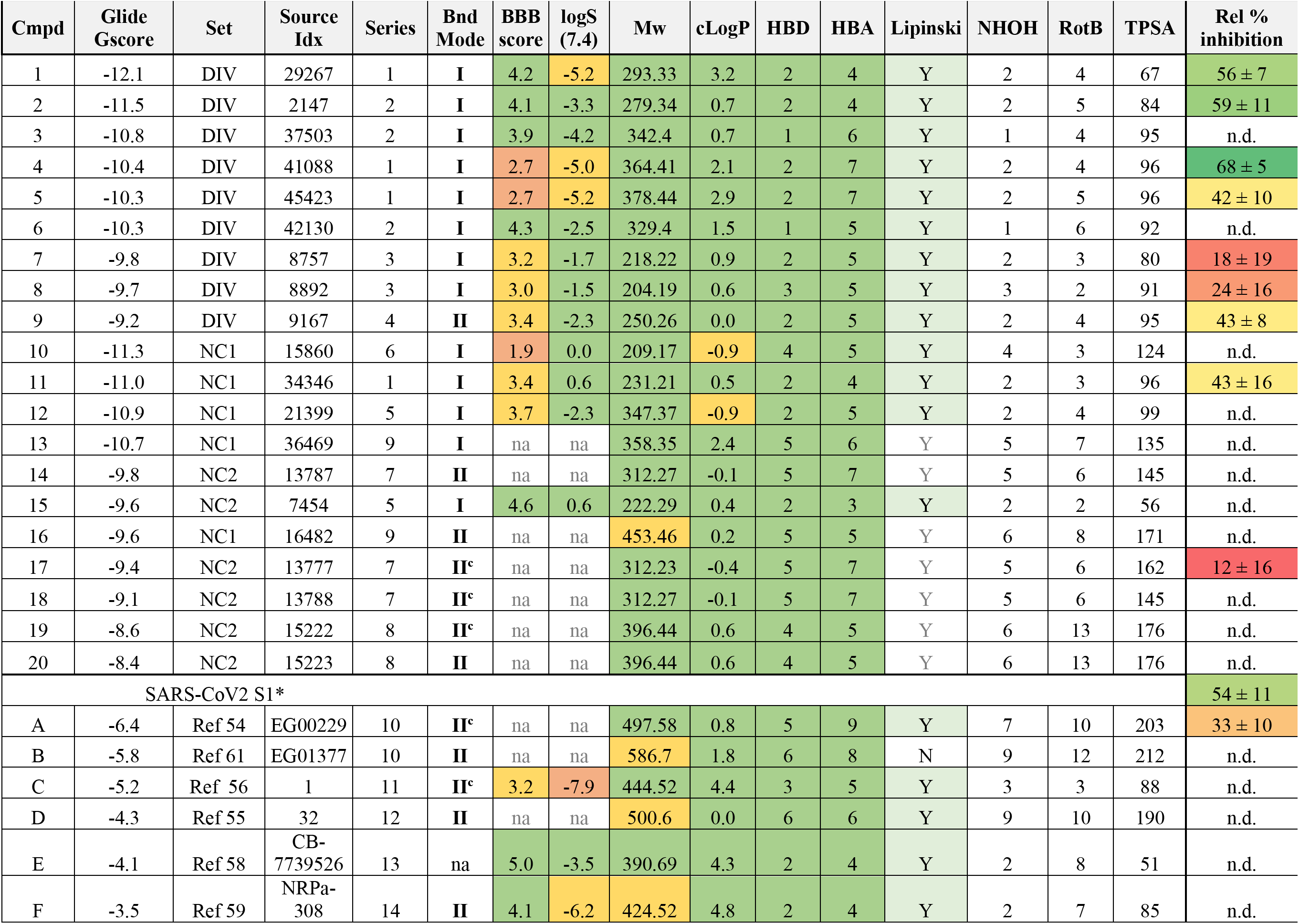
Hit and reference compounds with annotations of series assignment, binding Mode, origin, and calculated or experimental data. Compound ranking (Glide Gscore (kcal/mol)) and calculated properties of the top twenty hits with added reference compounds (A-F). Set: screening library;Source_Idx: reference to internal ID in the screening set; Series: chemotype assignment to hit series 1-9 and known compound series 10-14; Bnd Mode: binding Mode I or II (^c^ denotes constrained docking); BBB score: score for compound probability of having CNS exposure; LogS(7.4) predicted solubility (M) at pH 7.4; Mw molecular weight (Da); cLogP predicted lipophilicity coefficient in octanol/water; HBD number of hydrogen-bond donors; HBA number of hydrogen bond acceptors; Lipinski - a binary (Y/N) assignment of complying with Lipinski rule-of-5; NHOH number of polar NH and OH hydrogens; RotB number of rotatable bonds; TPSA total polar surface area (Å^2^); na denotes not applicable; n.d. indicates value not determined. All properties were calculated using RDKit. Relative % inhibition (ELISA VEGF-NRP-1 assay – see Methods section) with standard error of the mean (n=11).\

Since one objective of inhibiting the VEGF-A/NRP-1 interaction is disruption of pain signals, the full therapeutic effect will require drug exposure in the central nervous system (CNS). Correspondingly, the optimization strategy for these compounds will include modifications beneficial for crossing the blood-brain barrier (BBB), for example, decreasing the number of hydrogen bond donors and polarity. To supplement traditional medicinal chemistry approaches to increasing CNS exposure, we implemented the BBB score algorithm of Weaver et al.^67^ as one of the optimization parameters. Values of the BBB score in the range of 4 to 6 correctly predict 90% of CNS drugs^67^. While the brain/plasma ratio will be experimentally assessed for series representatives, optimization will be additionally guided by surrogate estimates of passive diffusion (PAMPA) and assessment of efflux and transporter proteins using assays in MDCK cell lines. Overall, predicted physico-chemical properties of all small molecule hit series (series 1-6) fall within ranges of lead-like and/or drug-like molecules (**Table 1**). Moreover, several representatives of series 1 and 2 are expected to have acceptable CNS exposure.

### Series analysis and docked binding modes

After initial ranking and selection of the top 20 hits, we proceeded to a more detailed analysis of structural and chemical features of the compounds. Chemical structures of all twenty hits and one virtual SAR analog are shown in **Figure 2**. Hits are grouped by common core motifs and molecular fragments that engage in productive hydrogen bond (HB) and alkyl/aryl π-contacts within the binding pocket. Inspection of aligned 2-D structures makes it apparent that the 2(1H)-pyridone core of structure **15** (highlighted in **Figure 2**) is the minimal motif of all small molecule hits (discussed further below). This core binds near the top of a central hydrophobic box formed by residues Tyr297, Tyr253, Trp301, and the methyl group of Thr316 (**Figure 3**). As expected, all of the hits bind within this box with the aryl or alkyl (e.g. isobutyl) groups engaged in hydrophobic interactions with these residues. Indeed, it is these hydrophobic interactions that are drivers of the overall binding affinity as judged by the Glide XP Gscore.

**Figure 2.**
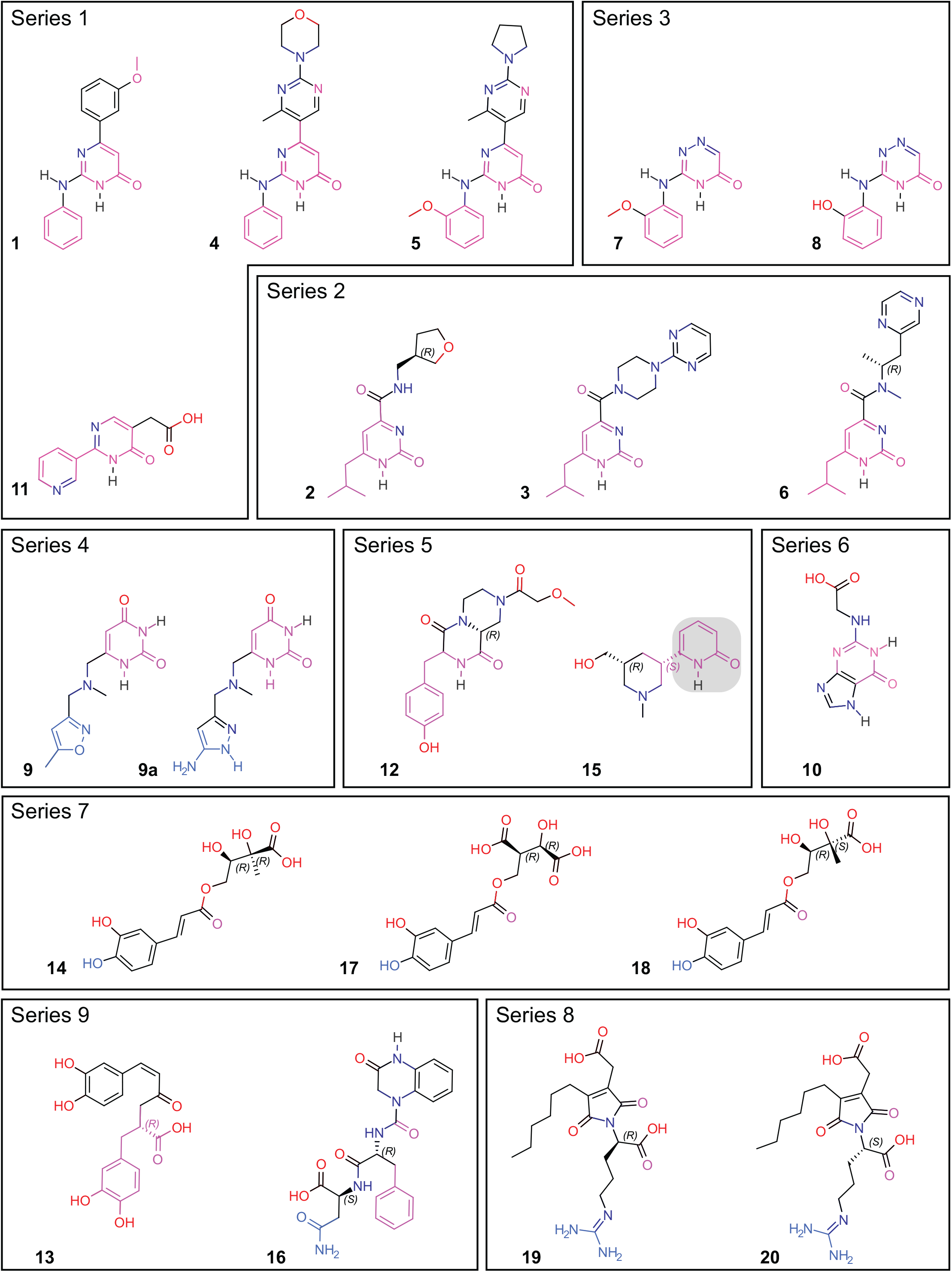
Chemical structures of the top 20 hits. Compounds are grouped by common core motif, shown in magenta. Molecules that adopt binding Mode II (**9, 9a, 14, 17, 18, 16, 19, 20**) have the atoms that form potential hydrogen bonds with Asp320, E319, or G318 colored in blue. The common 2(1H)-pyridone core is highlighted with a gray box in structure **15**.

**Figure 3.**
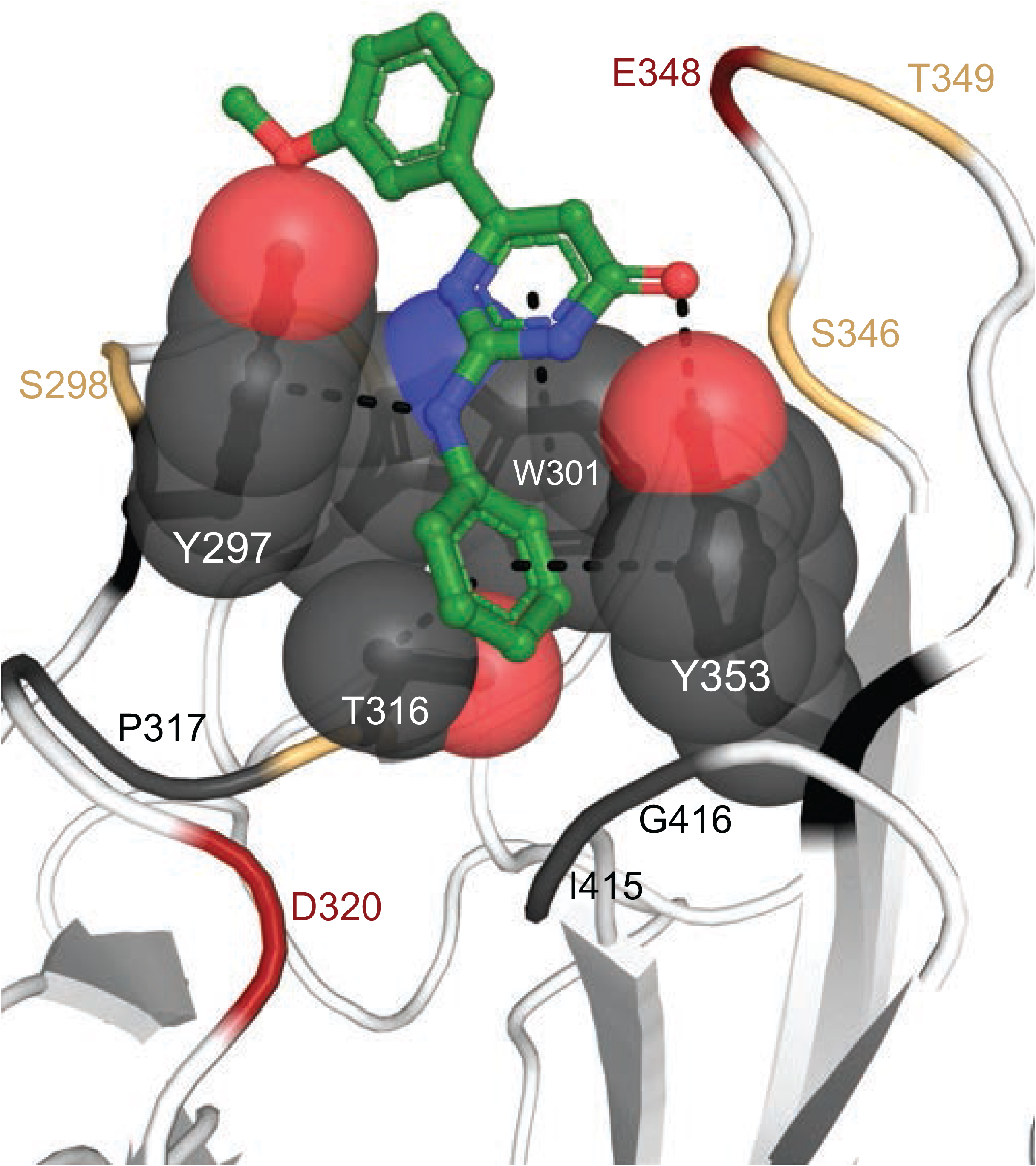
NRP-1 hydrophobic box. The hydrophobic groove formed by residues Tyr297, Tyr253, Trp301, and Thr316 are shown as gray spheres. Compound **1** is shown as green sticks and contacts with hydrophobic box residues within 5 Å shown as dashes. Remaining binding site residues colored red if acidic, yellow if polar, and gray if hydrophobic.

From this central position, the hits extend out of the aryl box in two general binding modes, which we refer to as Mode I (**Figure 4A**) and Mode II (**Figure 4B**). Interestingly, it is the core lactam carbonyl group in compounds **1-12, 15, 19-20** that makes potential hydrogen bonds with the hydroxyl groups of Thr349 and Tyr353 (**Figure 4C, D)** whereas in **14, 16**-**18** the carbonyl in the carboxylic group make these contacts, and in **13, 19, 20** the carboxyl makes contacts with S346, T349, and Y353 (**Figure 4E)**. Thus, all of compounds are able to partially mimic the terminal carboxyl contacts that are considered to be critical for anchoring the terminal Arg of CendR peptides^12, 31, 38-42, 44-53^ (**Figure 1C**) or known small molecule mimetics and inhibitors that contain a terminal guanidyl from Arg moiety and carboxylic group^54, 61, 62^.

**Figure 4.**
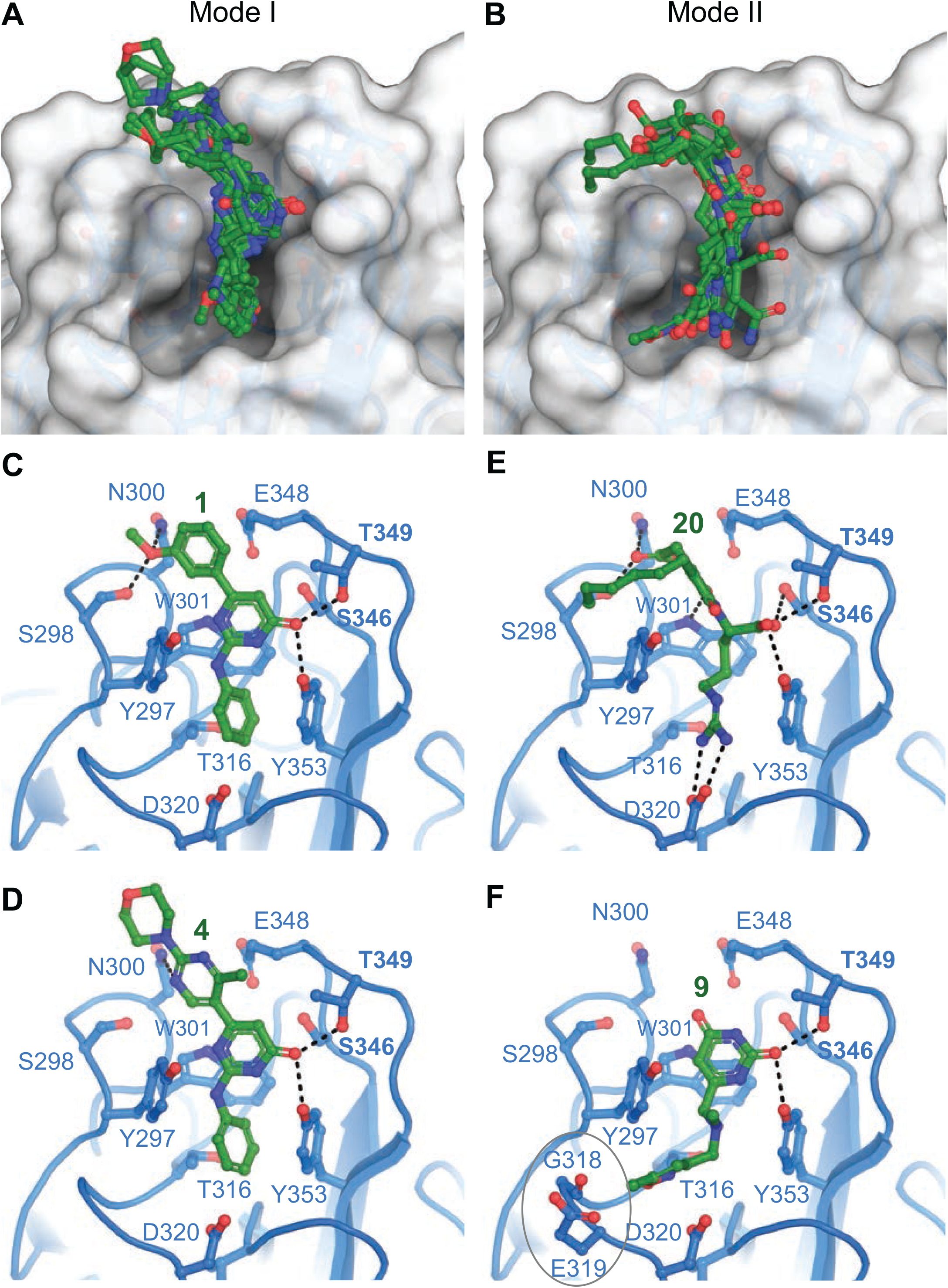
Hit compound poses in the NRP-1 b1-domain pocket. **A**. Overlay of hits that adopt binding Mode I. **B**. Overlay of hits that adopt binding Mode II. Polar interactions for representative compounds are shown as dashes for Mode I in **C**. (compound **1**) and **D**. (compound **4**) and for Mode II in **E**. (compound **20**) and **F**. (compound **9**).

All of the hit molecules occupying Mode I are synthetic compound chemotypes, including **11, 12**, and **15** from the NC1 and NC2 libraries, as noted above. Functional groups in **1, 2, 4, 5, 10, 12, 13** are involved in one or more potential hydrogen bond contacts with polar side chains Ser298 and Asn300 (**Figure 4C, D**). No other heteroatoms nor N-H hydrogen in pyrimidone (**1, 4, 5, 10, 11**), 2-pyrimidone (**2, 3, 6**), or 4H-1,2,4-triazin-5-one (**7, 8**) scaffolds are seemingly engaged in productive binding. Such “silent” polar sites provide the opportunity for replacement and optimization of ADME properties (e.g., oral absorption, systemic/CNS distribution) of these compounds. Interestingly, upon detailed analysis of hydrogen-bond patterns, we noticed that series **1, 3** but also **2-6** feature H-donor/H-acceptor topology of kinase inhibitors, including the presence of hydrophobic residues found in hinge-binding ATP mimics. Such features warrant the inclusion of kinase selectivity panels in the optimization stage. While kinase activity might be a feature to optimize out, our validation experiments support that binding of compounds from our series to the CendR site on NRP-1 (discussed below).

Mode II, with the exception of compound **9**, features skeletons of natural compounds. The functional groups in molecules **14** and **16-20** possess extensions toward the base of the pocket that form ionic or hydrogen bond contacts to residue Asp320 (**Figure 4E**). Furthermore, compound **9** extends toward the open region of the binding pocket bordered by Gly318 and Glu319 (**Figure 4F**). In preliminary SAR for compound **9**, we found that augmenting these interactions by the replacement of 5-methylisoxazole with 5-aminopyrazole (**9a, Figure 2**) led to an improvement in the Glide XP Gscore of 1.4 kcal/mol. Exploration of the interaction patterns observed in both binding modes is expected to improve binding affinity and compound selectivity.

We note that molecules **13** and **16**, while having the key pharmacophores present, had their geometry altered during the ligand preparation, likely a result of missing or incorrect chiral information in the COCONUT library, a known potential issue^63^. The hydroxycinnamyl group in **13** is present in the less stable Z-conformation and the chiral center at phenylalanine in **16** has inverted to (R)-configuration. Such alterations made both analogs unavailable from commercial sources of natural products. However, due to the availability of both precursors, derivatives **13** and **16** can be synthesized. Finally, the last two compounds (**19, 20**) are ionic, moderately reactive compounds which are not considered to be drug-like. Nevertheless, since both match the key features of CendR peptides (N-acylarginine), they provide valuable points for SAR.

### Comparison to known small molecules

To enable comparison with small molecules reported by others we docked and calculated physico-chemical properties of seven compounds that also target the NRP-1 CendR site^54-56, 58-61^. Based on chemotypes, we assigned these six molecules into unique series **10-14** (**Figure 5**). These molecules all exhibited lower docking scores than our hits (**Table 1**). Moreover, several of compounds **A-F** feature functional groups known for contributing to suboptimal physico-chemical and ADME properties, such as solubility, low intestinal absorption, metal chelation, and lability in liver microsomes or hepatocytes (**Table 1**). Docking poses for all compounds except **E** adopted Mode II binding.

**Figure 5.**
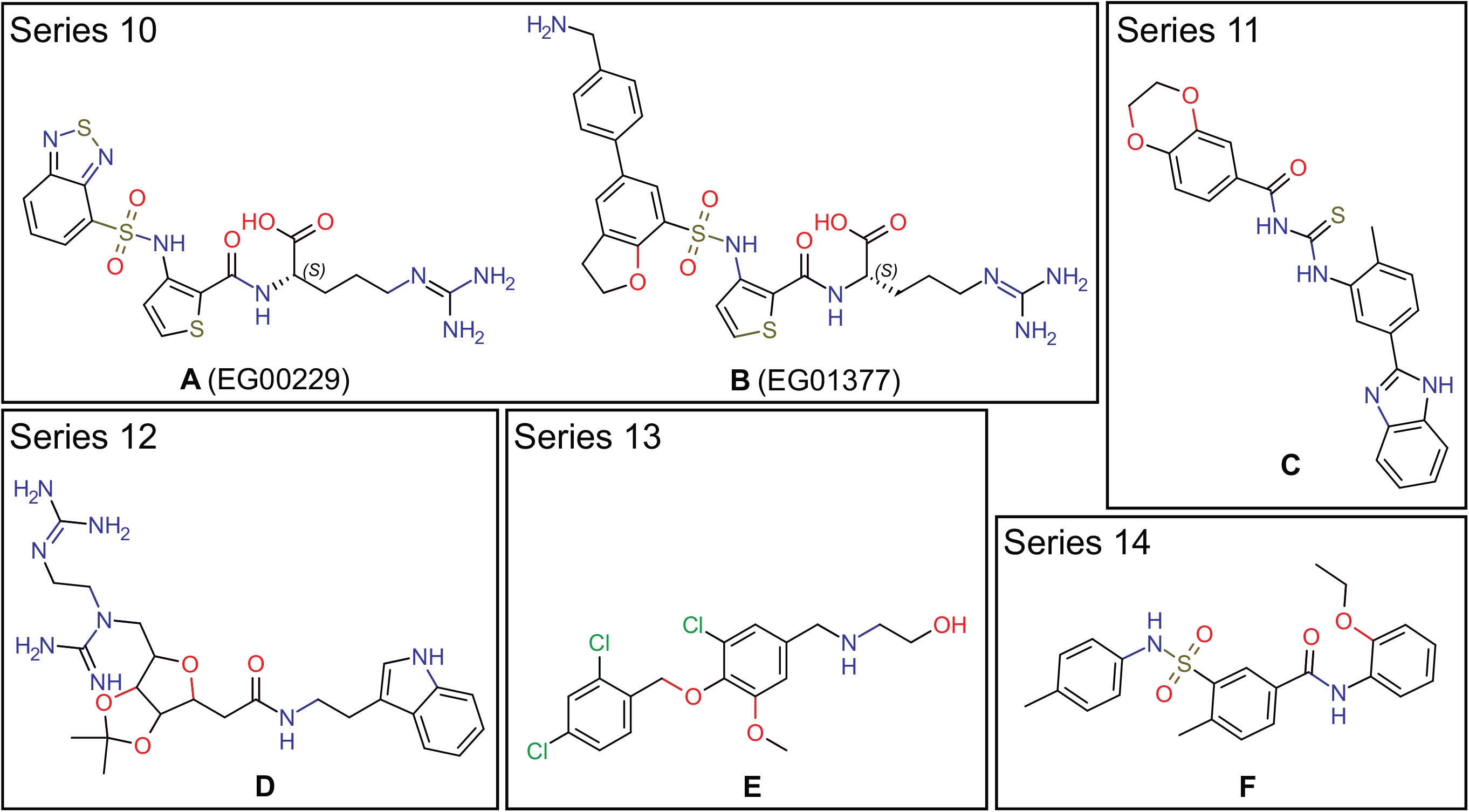
Structures of selected NRP-1 targeting compounds. Compound A (EG00229) from Ref^54^, compound B (EG01377) from Ref^61^, compond C is “compound 1” from Ref^56^, compound D is “bis-guanidinylated compound 32” from Ref^55^, compound E is ChemBridge ID: 7739526 from Ref^58^, and compound F is “NRPa-308” from Ref^59^.

### Validation of selected hits

We evaluated the ability of hit compounds to interfere with the NRP-1/VEGF-A interaction using an enzyme linked immunosorbent assay (ELISA). We coated plates with the extracellular domain of human NRP-1 (containing the a1a2 and the b1b2 regions) and added a selection of our compounds (based on SAR and commercial availability) to disrupt the NRP-1/VEGF-A interaction. We found that 3 hit compounds blocked more than 50% of VEGF-A binding to NRP-1. This level of inhibition mimicked that we observed with SARS-CoV-2 Spike protein (**Table 1**). In comparison, EG00229’s level of inhibition was outmatched by 6 of the 9 compounds we screened in this assay (**Table 1**). Thus, three of the synthetic compounds significantly inhibited the interaction between VEGF-A and NRP-1, confirming that they compete for binding to the CendR site (**Table 1**).

Compounds for biochemical evaluation were selected such that each structural feature was complementary to the overall SAR. Relative inhibition in the ELISA is reported in **Table 1**. Unfortunately, several compounds (**12, 14, 15, 18, 19, 20**) were not commercially available. Since many of those compounds can be synthesized in two to five steps, we intend to make essential representatives during future SAR optimization.

Inspection of docking scores and relative inhibition data in **Table 1** reveals a correlation between predictions and ELISA data. Most importantly, the binding hypotheses were confirmed as compounds with pyridone and pyrimidone cores, when appropriately decorated, were found to disrupt the VEGF-A/NRP-1 interaction more effectively than EG00229 and to a similar extent as SARS-CoV-2 Spike protein. As could be expected, interference was greater for compounds with additional functional groups (**1, 2, 4, 5** vs. **7, 8, 11**). Such a trend is important for the design of new analogs that will expand on underexplored scaffolds (**Series 3**, compound **11**). As discussed later, we intend to improve binding and inhibition by borrowing structural features from Modes I and II (e.g., compound **9a**).

Next, we set out to test the compounds for their capacity to inhibit the activation of the VEGF-A pathway. VEGF-A binding to the dimeric complex of its receptor VEGFR2 and co-receptor NRP-1 triggers phosphorylation of the VEGFR2 cytoplasmic domain at Y1175 (**Figure 6A**). Using an in-cell Western assay, we tested the compounds for their ability to inhibit increased phosphorylation of VEGFR2 by VEGF-A. In this assay, VEGF-A doubled the level of VEGFR2 phosphorylation at Y1175 (**Figure 6B**) which could be blocked by SARS-CoV-2 Spike protein as well as by the reference compound EG00229 (**Figure 6C**). All 9 of our tested compounds significantly blocked the VEGF-A stimulated increased phosphorylation of VEGFR2 (**Figure 6**). In the absence of stimulation by VEGF-A, only one of the 9 compounds (**4**) showed inhibition of basal VEGFR2 phosphorylation. As mentioned above, several of our compounds do exhibit features that are consistent with known kinase inhibitors which will be addressed during optimization of these hits.

**Figure 6.**
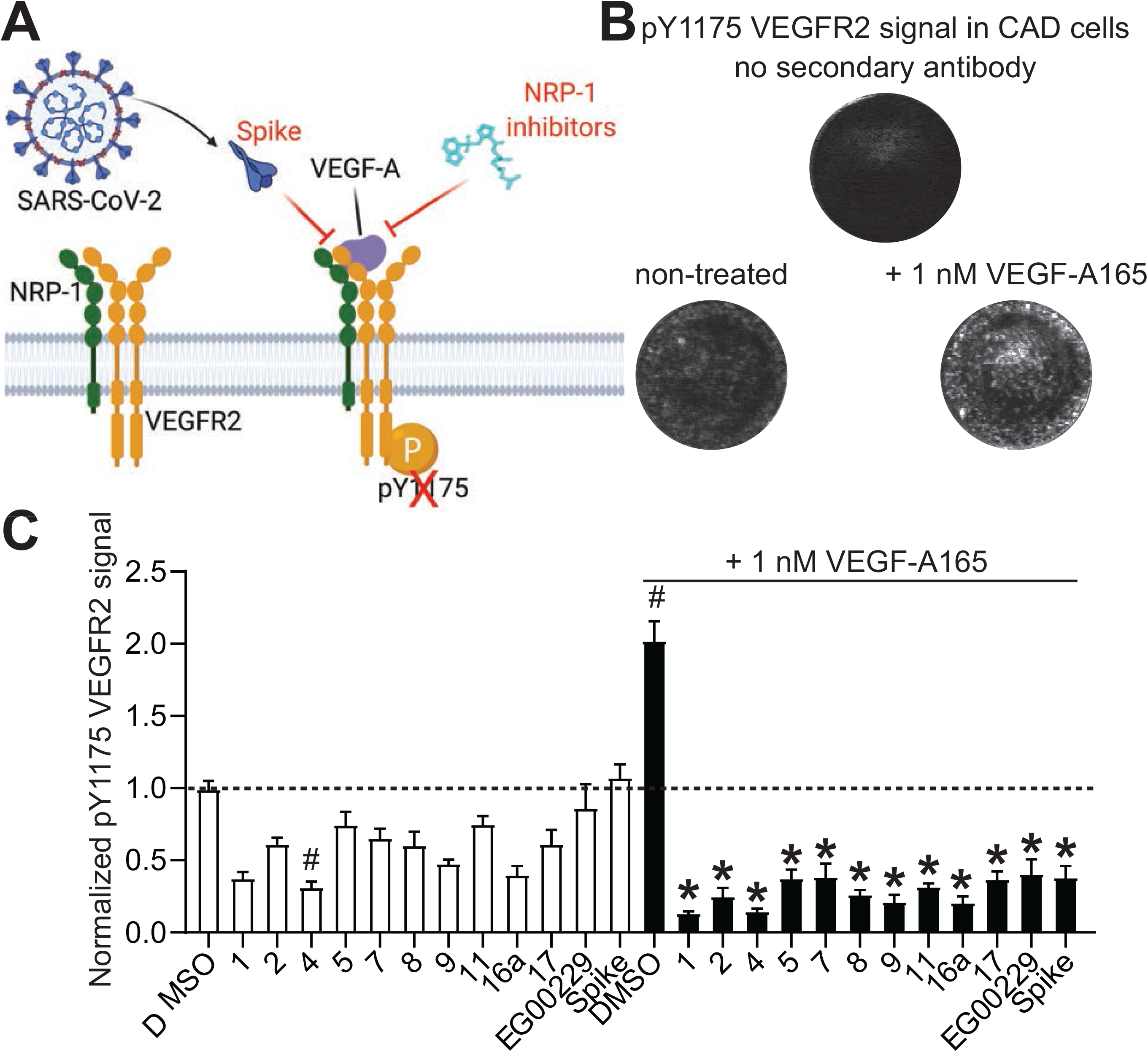
Screening of NRP-1/VEGF-A165 inhibitors by in cell western. (A) schematic of VEGF-A triggered phosphorylation of VEGF-R2. Screening of NRP-1/VEGF-A165 inhibitors by in-cell Western. Cathecholamine A differentiated (CAD) cells were grown in 96 well plates. Cells were treated with the NRP-1/VEGF-A165 inhibitors at 12.5 µM or SARS-CoV-2 Spike (100 nM) in combination with 1nM VEGF-A165 as indicated. Cells were stained for pY1175 VEGFR2 as a marker of the activation of the pathway by VEGF-A165. (B) representative micrographs showing the lack of signal in controls with omission of the secondary antibody. Phosphorylated VEGFR2 was increased by the addition of 1 nM VEGF-A165 on the cells. (C) Bar graph showing the levels of pY1175 VEGFR2 normalized to the quantity of cells in each well. # p<0.05 compared to 0.1% DMSO (vehicle) treated cells without VEGF-A165. * p<0.05 compared to 0.1% DMSO + 1 nM VEGF-A165 treated cells, Kruskal-Wallis test with Dunn’s multiple test correction (n=7 replicates per condition). Data was analyzed by a repeated measures one-way analysis of variance (post hoc: Dunnett’s), *p<0.05.

Finally, we screened the compounds for antiviral activity using a GFP-expressing vesicular stomatitis virus (VSV) recombinant protein, encoding the SARS-CoV-2 spike protein rather than the native envelope glycoprotein^68^. This VSV-eGFP-SARS-CoV-2 mimics SARS-CoV-2 and is a convenient BSL2 platform to assess SARS-CoV-2 Spike-dependency. Vero-E6-TMPRSS2 cells, which overexpress the transmembrane serine protease 2 (TMPRSS2)^68^, were infected in the presence of individual compounds or DMSO vehicle. GFP fluorescence was measured 36 hours post infection by automated microscopy (**Figure 7A**). Two compounds (**1** and **5**) displayed >50% inhibition of VSV-eGFP-SARS-CoV-2 antiviral activity while another two compounds (**16a, 17**) demonstrated ∼15% inhibition (**Figure 7B**). Spike inhibited antiviral activity by ∼35% while the known NRP-1 inhibitor EG00229 was ineffective in this assay.

**Figure 7.**
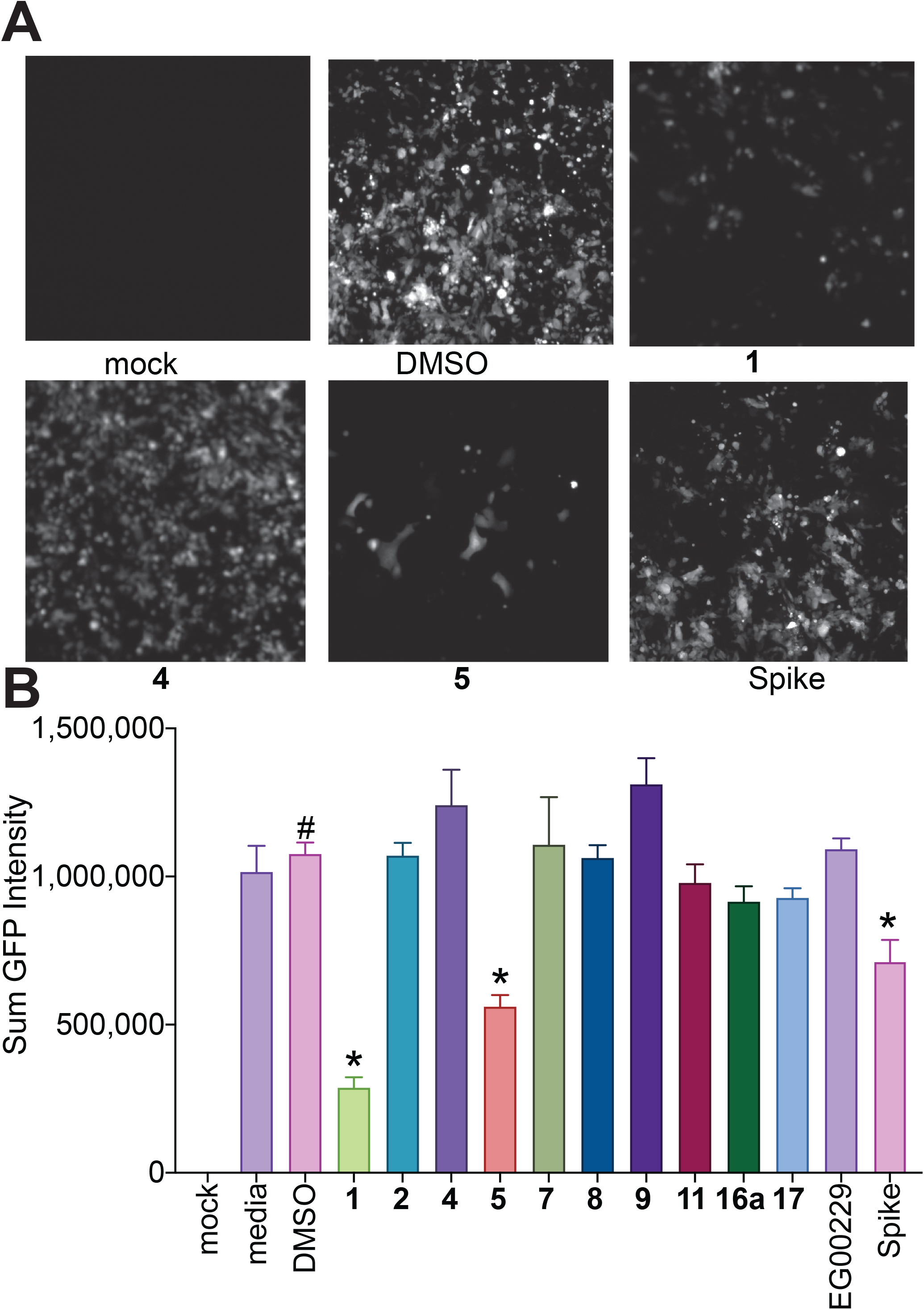
Screening for VSV-eGFP-SARS-CoV-2 inhibition. Compounds were screened at 25 μM for inhibition of VSV-eGFP-SARS-CoV-2 infection of Vero-E6-TMPRSS2 cells. Recombinant Spike S1 domain was included at 68 nM. Cells were infected for 36h prior to live cell automated microscopy and quantification of sum GFP fluorescence intensity, normalized to cell count by HCS CellMask Blue, was measured and for each well and plotted with Prism 6. Results are presented as mean intensity ± SEM, # P <0.05 vs. mock; *P < 0.05 versus DMSO (n=3 replicates). Data was analyzed by a one-way analysis of variance (post hoc: Dunnett’s), *p<0.05.

### Pharmacophore models

A pharmacophore model was derived from the identified ligands, considering both steric and electronic requirements (**Figure 8**). The most critical features are the aromatic rings A1, A2, and the hydrogen bond acceptor HBA. This HBA is typically a carbonyl oxygen engaged in contacts with the hydroxyl groups of Tyr353 and Thr349. The aryl group in A1 directs the carbonyl oxygen of the HBA toward those residues. Alternatively, A1 can be presented in an edge-to-face contact with Tyr297. There are two additional acceptor sites on the opposite side of the A1 ring, relative to acceptor HBA. These can form hydrogen bonds with the side chain of the Asn300 amide or the Ser298 hydroxyl. The area between the HBA and these two additional acceptors (where the label A1 is located) is expected to accommodate a structural molecule of water^9, 13, 60, 61^. This water has been proposed to be important in ligand binding as it may bridge interactions to Trp301^60^. However, our screens were conducted in the absence of water molecules. Nevertheless, we observe that polar groups in several analogs occupy the position of this structural water and get involved in the corresponding hydrogen bond network, suggesting they could displace it. Aromatic ring A2 is sandwiched in the hydrophobic box formed by residues Y353, W301, Y297, and the methyl group of T316. Binding mode II includes A1/HBA, A2, and features an expansion toward polar residues (E319, D320) and additional stabilizing contacts in the lower part of the pocket. It is this area (donor, donor, acceptor) which accepts the guanidino group of the CendR Arg and contains the open lower left pocket seen in Mode I (Figure 4A). By connecting all important pharmacophore features, hybrid synthetic molecules can be envisioned that will merge binding modes I and II and extend into the lower left to fully occupy the available binding pocket.

**Figure 8.**
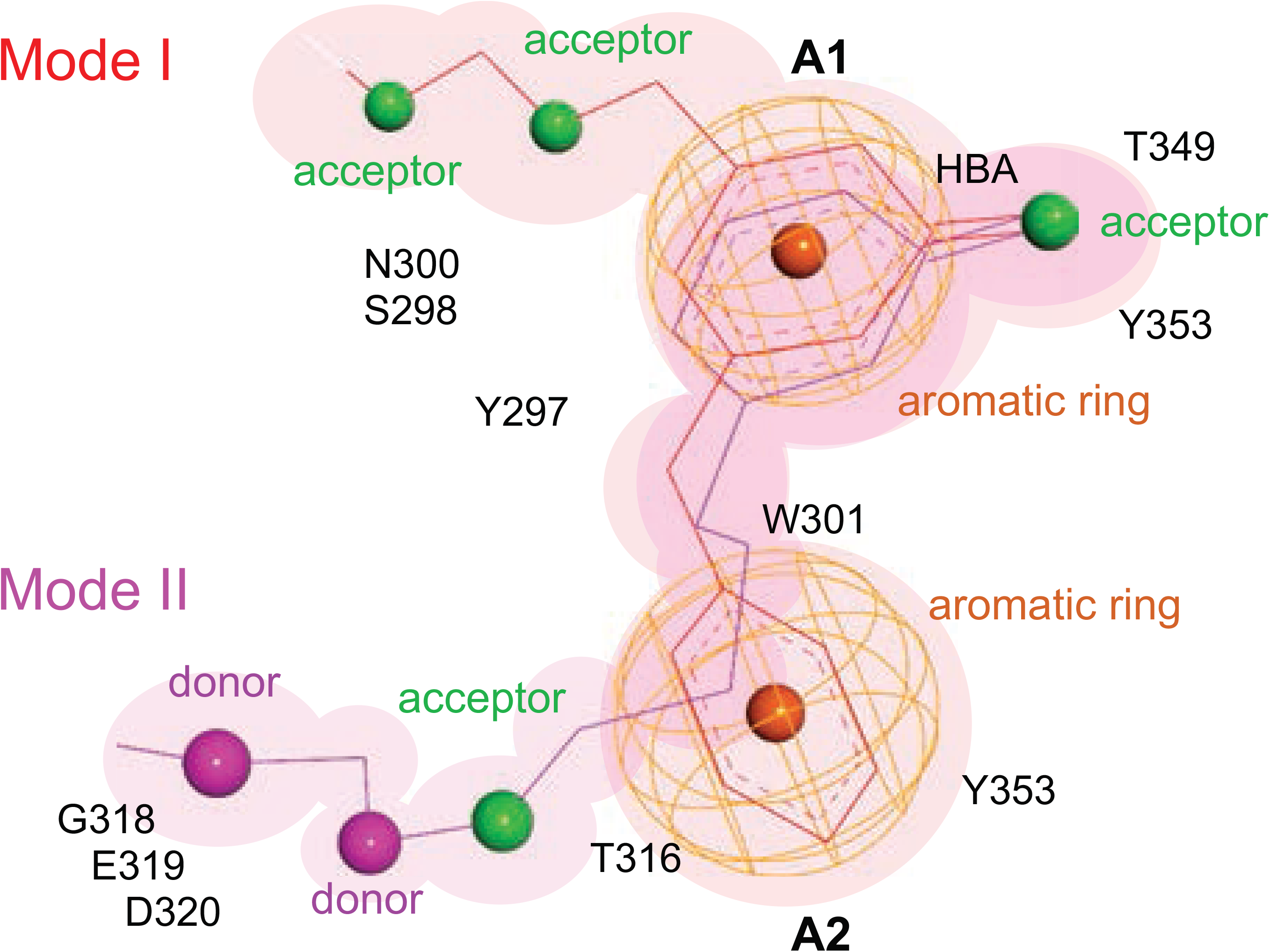
Schematics of pharmacophore models. Thin lines indicate bond distances between pharmacophores while the color indicates the mode of binding (Mode I in red, Mode II in magenta). Pharmacophore features are shown in color (acceptors in green, donors in magenta, and aromatic rings as orange mesh). The neighboring or contact protein residues are listed.

## Conclusions

From a virtual screen of nearly 0.5 M compounds, we identified nine chemical series comprising small molecules and natural products that occupy the CendR binding site on NRP-1. All compounds identified in our series fall within ranges of lead-like and/or drug-like molecules which enhances their potential for efficacy in *in vitro* and *in vivo* assays. This *in silico* effort allowed us to identify two modes of binding within the CendR pocket. To guide future drug discovery efforts, we propose a hybrid pharmacophore model that will enable design of small molecules that will maximize the pocket occupancy. Two validation experiments confirmed that a subset of our hits compete with binding of VEGF-A and interfere with VEGF-A induced phosphorylation of VEGFR2, supporting direct binding to the CendR site on the b1 domain. Two CendR-blocking compounds inhibited Spike-dependent infection of VSV-eGFP-SARS-CoV-2 and may have potential for further development, although additional studies are needed to understand their antiviral mechanisms and involvement of NRP-1 and ACE-2 receptors. To guide future drug discovery efforts, we propose a hybrid pharmacophore model that will enable design of small molecules that will maximize pocket occupancy and contacts. Since the VEGF-A/NRP-1 signaling pathway participates in multiple pathologies including neuropathic pain and cancer, our series of lead compounds represent a first step in a renewed effort to develop small molecule inhibitors for the treatment of these diseases.

Finally, we mention one additional interesting aspect of this system that is still being explored. Heparin, the widely used anticoagulant drug is routinely used for hospitalized SARS-CoV-2 patients to lower the probability of blood clothing and embolism^69^. It is also known that heparin prevents infection by a range of viruses^70^ and was recently reported to inhibit invasion by SARS-CoV-2 in a cell-based assay^71^. Heparin is a required co-receptor for VEGF-A signaling^72^ and NRP-1 also binds heparin, mainly through the b1b2 domain, through sites distal to the CendR pocket^73^. This raises the possibility that the interaction of SARS-CoV-2 Spike/RBD with NRP-1 is facilitated by heparin and invites speculation of a potential synergistic effect of heparin and NRP-1 inhibitors as an efficacious drug combination to prevent viral entry.

## Methods

### Preparation of receptor protein and grid for virtual screening

Preparation and virtual screening steps were conducted using Schrödinger Release 2019-3 (Schrödinger, LLC, New York, NY, 2020). The highest resolution structure of the NRP-1 b1 domain was selected for docking (PDB ID: 6fmc)^61^. This structure was prepared using the Protein Preparation Wizard^74^ to remove all water molecules and alternate conformations, add and refine hydrogen atoms, and conduct restrained minimization (OPLS3e force field, convergence to 0.30 Å). There were no residues with alternate conformations within the binding pocket. A 20×20×20 Å grid box was centered on the co-crystallized inhibitor EG01377 to target the VEGF-A_165_ site. An optional, symmetric constraint was generated that required hit compounds to form a hydrogen bond to the side-chain of Asp 320.

### Screening libraries

The synthetic compound library (DIV) was obtained by combining ChemBridge Diversity Core and Express sets of drug-like compounds. These were prepared for screening in LigPrep using the OPLS3e force field, neutral ionization, desalting, and tautomer generation. If specified, chirality centers were maintained, otherwise up to three chiral variations were generated per atom and ligand. This library contained a total of 210,677 compounds (293,251 conformers). The COlleCtion of Open NatUral producTs (COCONUT) set of open-access natural compounds^63^ was downloaded from https://zenodo.org/record/3778405#.Xs1D6mhKiUk (on 5/26/20) and prefiltered by excluding compounds with molecular weight ≥ 500 Da and alogP ≥ 5. LigPrep settings were the same as for the DIV set and the resulting library (NC1) consisted of 257,166 natural compounds (50,686 conformers). The smaller natural compound library (NC2) library was a curated set of 20,088 natural compounds (23,846 conformers) originally obtained from ZINC15^64^. The NC2 library had some overlap with the NC1 library, but nevertheless produced useful results.

### Virtual screening and scoring

Virtual screens were run for each library against the VEGF-A_165_ binding site of NRP-1 using the Glide virtual screening workflow (Schrödinger, LLC, New York, NY, 2020)^75^. For the DIV and NC1 libraries, the default docking settings were accepted, with 10% of compounds at each stage (high-throughput virtual screen, standard precision docking, extra precision docking) resulting in 293 hits for the DIV library and 550 hits for the NC1 library. Because the NC2 library was smaller, it was set to retain 25%, 20%, 15% of the hits at each stage, resulting in 152 hits. The virtual screens were first run without and then with the use of the Asp 320 constraint, but only the constrained hits from NC2 were retained due to strained conformations and lower docking scores for the DIV and NC1 screens. Thus, a total number of 1,147 virtual hit compounds were obtained from 4 screens.

### Docking of known NRP-1 targeting compounds

Representatives of known compound series^54-56, 58, 59, 61^ were prepared for screening in LigPrep using the OPLS3e force field, neutral ionization, desalting, and tautomer generation. Docking was run against the VEGF-A_165_ binding site of NRP-1 using Glide XP (Schrödinger, LLC, New York, NY, 2020)^75^.

### Compound property calculations

The following physico-chemical properties were calculated using RDKit^76^: molecular weight in Daltons (Mw, Da), partition coefficient (clogP), number of hydrogen-bond donors and acceptors (HB-D, HB-A), total number of N-H and OH groups (NHOH), number of rotatable bonds (RotB), and total polar surface area (TPSA, Å^2^). To estimate compound solubility, calculated logS (M) at pH=7.4 was obtained using ChemAxon Aqueous solubility module^77^.

### ELISA□based NRP1-VEGF-A165 protein binding assay

Plates (96□well, Nunc Maxisorp; Thermo Fisher Scientific, Waltham, MA, USA) were coated with human Neuropilin-1-Fc (10 ng per well, Cat# 50-101-8343, Fisher, Hampton, NH) and incubated at room temperature overnight. The following day, the plates were washed and blocked with 3% BSA in PBS to minimize non□specific adsorptive binding to the plates. SARS-CoV2 Spike protein (100 nM, S1 domain aa16-685, Cat# Z03485, Genscript, Piscataway, NJ), EG00229 (Cat#6986, Tocris) or the indicated compounds were added at 12.5µM and incubated for 30 min at room temperature prior to adding biotinylated human VEGF-A165 (Cat#BT293, R&D systems) at 10 nM. As a negative control, some wells received PBS containing 3% BSA. The plates were incubated at room temperature with shaking for 1 h. Next, the plates were washed with PBS to eliminate unbound protein. Bound biotinylated VEGF was detected by streptavidin-HRP (Cat#016-030-084, Jackson immunoResearch). Tetramethylbenzidine (Cat#DY999, R&D Systems, St. Louis, MO) was used as the colorimetric substrate. The optical density of each well was determined immediately, using a microplate reader (Multiskan Ascent; Thermo Fisher Scientific) set to 450 nm with a correction wavelength of 570 nm. Data were normalized to the background and to the signal detected for VEGF-A165 alone.

### In cell western for detecting inhibition of VEGFR2 activation by VEGF-A165

Mouse neuron derived Cathecholamine A differentiated CAD (ECACC, Cat# 08100805) were grown in standard cell culture conditions, 37 °C in 5% CO2. All media was supplemented with 10% fetal bovine serum (Hyclone) and 1% penicillin/streptomycin sulfate from 10,000 µg/ml stock. CAD cells were maintained in DMEM/F12 media. Cells were plated in a 96 well plate and left overnight. The next day, indicated compounds (at 12.5µM), or SARS-CoV-2 Spike (100 nM, S1 domain) were added in CAD cell complete media supplemented with 1 nM of mouse VEGF-A165 (Cat# RP8672, Invitrogen) and left at 37°C for 1 hour. Media was removed and the cells rinsed three times with PBS before fixation using ice cold methanol (5 min). Methanol was removed and cells were left to dry completely at room temperature. Anti-VEGFR2 pY1175 was used to detect the activation of the pathway triggered by VEGF-A165 in the cells. The antibody was added in PBS containing 3% BSA and left overnight at room temperature. The cells were washed three times with PBS and then incubated with Alexa Fluor® 790 AffiniPure Goat Anti-Rabbit IgG (Cat# 111-655-144, Jackson immunoResearch) in PBS, 3% BSA for 1 hour at room temperature. Cells were washed three times with PBS and stained with DAPI. Plates were imaged on an Azure Sapphire apparatus. Wells that did not receive the primary antibody were used a negative control. The signal was normalized to the cell load in each well (using DAPI) and to control wells not treated with VEGF-A165.

### Cell and viral culture

Vero-E6-TMPRSS2 cells were cultured in high glucose DMEM supplemented with 10% FBS and 1% penicillin/streptomycin. Cells were supplemented with 20 μg/mL blasticidin (Invivogen, ant-bl-1) to maintain stable expression of TMPRSS2 during routine culture. Cells were maintained at 37°C with 5% CO_2_ and passaged every 2-3 days. These African green monkey Early kidney cells express NRP-1 which is 99.3% similar within its b1 domain to the human NRP-1. Homology models (not shown) also reveal no differences in NRP-1 passage between these species. infectious VSV-eGFP-SARS-CoV-2 stock was a generous gift from Sean P.J. Whelan (Washington University, St. Louis, MO, USA). VSV-eGFP-SARS-CoV-2 was passaged once by infecting Vero-E6-TMPRSS2 cells at MOI = 0.01 in DMEM + 2% FBS and 1% penicillin/streptomycin for 72 hr at 34°C. Cell-free supernatant was collected and concentrated 10-fold through Amicon-Ultra 100 kDa MWCO spin filter units (Millipore UFC905008) prior to aliquoting and storage at -80°C. Titer was determined to be approximately 1×10^7^ PFU/mL as determined by median tissue culture infectious dose (TCID_50_) assay on Vero-E6-TMPRSS2 cells.

### Screening compounds for VSV-eGFP-SARS-CoV-2 inhibition

African green monkey kidney (Vero)-E6-TMPRSS2 cells were plated at 15,000 cells per well in black/clear bottom 96 well tissue culture plates (Thermo Fisher Scientific 165305). The next day cells were infected ± 0.001% DMSO, 25 μM compounds, or 68 nM recombinant Spike protein, at an MOI of 0.05 in 100 μL DMEM + 10% FBS and 1% penicillin/streptomycin for 36 hr at 37°C prior to live cell fluorescent microscopy on a Nikon Eclipse Ti2 automated microscopy system with 4x objective and 488/532 nm filters. Sum GFP fluorescence intensity, normalized to cell count by HCS CellMask Blue (Thermo #H32720), was measured and for each well and plotted with Prism 6 (GraphPad Software). Significant differences were determined by RM one-way ANOVA followed by multiple comparisons test. An alpha of 0.05 was used to determine the statistical significance of the null-hypothesis.

## Acknowledgments

We thank Sean P.J. Whelan and Paul W. Rothlauf for generously sharing the VSV-eGFP-SARS-CoV-2 clone and the Vero-E6-TMPRSS2 cells.

## Funding

Supported by NINDS (NS098772 (R.K.)), NIDA (DA042852, R.K.), NIGMS (GM136853, S.K.C.), and a UA

COVID-19 seed grant (002196, S.K.C.)

## Author contributions

R.K. and A.M. developed the concept; S.P.-M. and M.P. conducted virtual screening, docking and analysis; A.M. performed the ELISA and in-cell western experiments and analyzed the data; S.K.C. performed the VSV-SARS-CoV-2 infections; C.R.C. performed microscopy and collected the VSV-SARS-CoV-2 infection data; C.A.T. and S.K.C. supervised and analyzed the infection experiments. All authors wrote and edited the manuscript; and R.K. supervised all aspects of this project. All authors had the opportunity to discuss results and comment on the manuscript;

## Competing interests

R. Khanna is the co-founder of Regulonix LLC, a company developing non-opioids drugs for chronic pain. In addition, R. Khanna has patents US10287334 and US10441586 issued to Regulonix LLC. The other authors declare no competing financial interests.

## Data and materials availability

All data is available in the main text, figures, and supplementary materials.

## Abbreviations

ACE-2: angiotensin converting enzyme-2
ADME: absorption, distribution, metabolism, and excretion
BBB: blood brain barrier
CNS: central nervous system
COCONUT: COlleCtion of Open NatUral producTs
COVID-19: coronavirus disease 2019
ELISA: enzyme linked immunosorbent assay
NC: natural compound
NRP-1: Neuropilin 1
PDB: protein data bank
Ro5: Lipinski rule of 5
SARS-CoV-2: severe acute respiratory syndrome coronavirus 2
VEGF-A: vascular endothelial growth factor-A
VSV: vesicular stomatitis virus

## Table captions

**Supplementary Table 1**. Physico-chemical properties and structures of top 100 hits, arranged by docking score.

